# Novel tau filament folds in individuals with *MAPT* mutations P301L and P301T

**DOI:** 10.1101/2024.08.15.608062

**Authors:** Manuel Schweighauser, Yang Shi, Alexey G. Murzin, Holly J. Garringer, Ruben Vidal, Jill R. Murrell, M. Elena Erro, Harro Seelaar, Isidro Ferrer, John C. van Swieten, Bernardino Ghetti, Sjors H.W. Scheres, Michel Goedert

## Abstract

Mutations in *MAPT*, the microtubule-associated protein tau gene, give rise to cases of frontotemporal dementia and parkinsonism linked to chromosome 17 (FTDP-17) with abundant filamentous tau inclusions in brain cells. Individuals with pathological *MAPT* variants exhibit behavioural changes, cognitive impairment and signs of parkinsonism. Missense mutations of residue P301, which are the most common *MAPT* mutations associated with FTDP-17, give rise to the assembly of mutant four-repeat tau into filamentous inclusions, in the absence of extracellular deposits. Here we report the cryo-EM structures of tau filaments from five individuals belonging to three unrelated families with mutation P301L and from one individual belonging to a family with mutation P301T. A novel three-lobed tau fold resembling the two-layered tau fold of Pick’s disease was present in all cases with the P301L tau mutation. Two different tau folds were found in the case with mutation P301T, the less abundant of which was a variant of the three-lobed fold. The major P301T tau fold was V-shaped, with partial similarity to the four-layered tau folds of corticobasal degeneration and argyrophilic grain disease. These findings suggest that FTDP-17 with mutations in P301 should be considered distinct inherited tauopathies and that model systems with these mutations should be used with caution in the study of sporadic tauopathies.

## Introduction

In the adult human brain, six tau isoforms are expressed from a single gene through alternative mRNA splicing [1]. These isoforms differ by the inclusion/exclusion of two exons near the N-terminus (exons 2 and 3) and a single exon near the C-terminus (exon 10) of the protein. Exon 10 encodes a repeat of 31 amino acids and its inclusion gives rise to three isoforms containing four repeats (4R). The other three isoforms lack exon 10 expression and have three repeats (3R). These repeats and adjoining sequences constitute the microtubule-binding domains of tau. Part of this sequence also forms the core of assembled tau in neurodegenerative diseases, indicating that the physiological function of microtubule binding and the pathological assembly into amyloid filaments are mutually exclusive.

Mutations in *MAPT* lead to the formation of filamentous inclusions that are made of either 3R, 4R or 3R+4R tau [2]. Mutations that cause the relative overproduction of wild type 3R or 4R tau result in the deposition of 3R tau with the Pick fold [3] or 4R tau with the argyrophilic grain disease (AGD) fold [4]. Filamentous inclusions of 3R+4R tau associated with missense mutations V337M and R406W in *MAPT* adopt the Alzheimer fold [5].

Mutations of residue P301 [P301L, P301S and P301T] [6–9] are the most common mutations in *MAPT* that are associated with FTDP-17. Most individuals with mutations P301L and P301S present clinically with behavioural-variant FTD, with or without parkinsonism [10–13]. Mutation P301T has been shown to cause a clinicopathological picture of type III globular glial tauopathy (GGT) [9,14]. Mutation P301L has also been shown to give rise to cases of GGT. All three mutations cause 4R tauopathies with abundant filamentous tau inclusions in nerve cells and glial cells. Only mutant tau is deposited in individuals with the P301L mutation [15,16]. *In vitro* experiments have shown that, of the *MAPT* mutations tested, P301L and P301S tau had the least potential to promote microtubule assembly and the greatest ability to form heparin-induced filaments [8,17–21]. It has been suggested that mutations of residue P301 destabilise local structure and expose the sequence ^306^VQIVYK^311^ (PHF6) [22], which is necessary for the assembly of tau into filaments [23]. Seeds of assembled recombinant P301L tau nucleate P301L, but not wild-type, tau [24]. P301L and P301S are also the most widely used *MAPT* mutations in transgenic mouse models of tauopathies, because their overexpression gives rise to robust phenotypes consisting of tau hyperphosphorylation, filament formation and neurodegeneration [25–28]. In these models, the assembly into tau filaments is necessary for neurodegeneration, because overexpression of human P301S tau without the PHF6 sequence does not result in neurodegeneration [29].

Heterozygous missense mutations that give rise to an amino acid substitution at position 301 result in approximately 75% of the total tau protein being wild-type and 25% being mutant. In addition, recombinant P301L and P301S tau proteins have been shown to have a partial loss of microtubule polymerisation function [8,17], implying that at most 10-15% of soluble tau will have lost its function. In view of this modest reduction, loss-of-function toxicity of tau is unlikely to be the cause of FTDP-17, contrary to what has been suggested [30]. Instead, the ordered assembly into filaments made of mutant tau is likely the gain-of-toxic function mechanism giving rise to FTDP-17 [31].

By cryo-EM, we reported different structures of tau filaments extracted from the brains of transgenic mice overexpressing human P301S tau downstream of the Thy1 or the prion protein promoter [32]. The cryo-EM structure of P301L tau filaments extracted from the brains of rTg4510 transgenic mice has also been reported [33]. Here we present the cryo-EM structures of tau filaments extracted from the cerebral cortex of individuals with mutations P301L and P301T in *MAPT*. New tau folds were observed for both mutations, suggesting that mutations of residue P301 in tau lead to diseases that are distinct from sporadic tauopathies.

## Results

### Structure of tau filaments from five cases of three unrelated families with mutation P301L in *MAPT*

We determined the structures of tau filaments from the cerebral cortex of five individuals belonging to three families with mutation P301L in *MAPT* (Figures 1 and 2). Parietal cortex was used for cases 1 and 2, and temporal cortex for cases 3, 4 and 5. All cases had only tau filaments made of an identical single protofilament, with resolutions of 2.8-3.9 Å (Figure 1a). For the sharpened map in Figure 2, we used tau filaments that were extracted from the parietal cortex of case 2, as this gave the highest resolution. According to the genomic analysis, the three families were unrelated, but the individuals from the Dutch family (cases 3, 4 and 5) were related. All five individuals had a C to T nucleotide substitution in the second position of codon 301 (CCG to CTG, on one allele), resulting in a P301L change. The clinicopathological diagnosis of all cases was behavioural-variant FTD. By immunoblotting of sarkosyl-insoluble fractions with BR133, RD3, anti-4R, BR134, AT8 and AT100, strong tau bands of 64 and 68 kDa were observed with all antibodies, except RD3, indicating the presence of hyperphosphorylated 4R, but not 3R, tau (Figure 1b). Assembled tau was truncated at the N-terminus, as judged by the presence of additional lower molecular weight tau bands with anti-4R, BR134, AT8 and AT100, but not with BR133. By immunohistochemistry, abundant inclusions of hyperphosphorylated 4R tau were present in nerve cells and glial cells, chiefly astrocytes (Extended Data Figures 1-4). They were Gallyas-Braak silver-positive. Globular glial tau inclusions were not observed.

**Figure 1.**
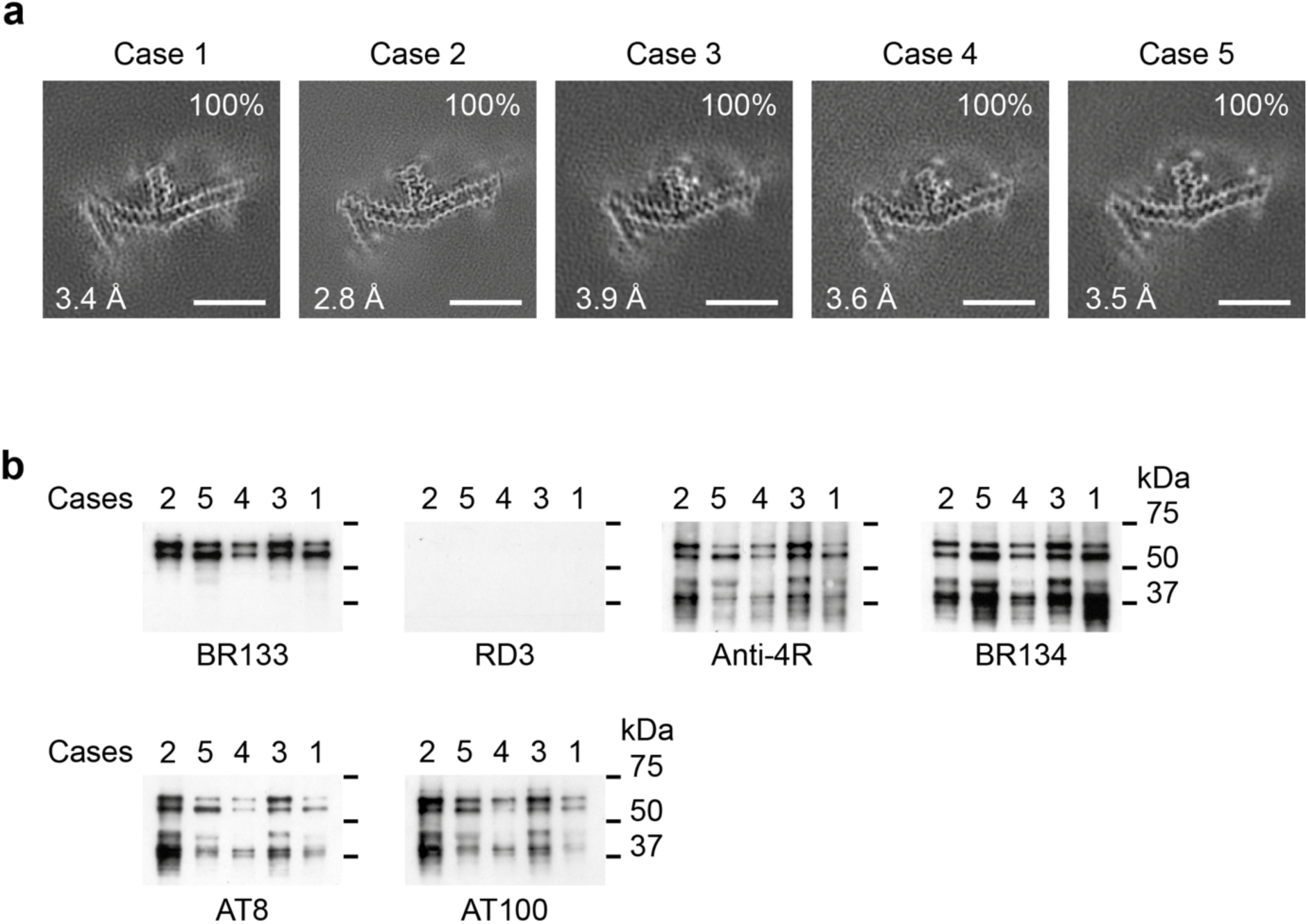
P301L mutation in *MAPT*: Cryo-EM cross-sections of tau filaments and immunoblotting. a, Cross-sections through the cryo-EM reconstructions, perpendicular to the helical axis and with a projected thickness of approximately one rung, are shown for the parietal cortex from cases 1 and 2, and the temporal cortex from cases 3, 4 and 5. Resolutions (in Å) and percentages of filament types are indicated at the bottom left and top right, respectively. All filaments had the three-lobed fold of assembled P301L tau. Scale bars, 10 nm. b, Immunoblotting of sarkosyl-insoluble tau from the parietal cortex of cases 1,2 and from the temporal cortex of cases 3, 4 and 5. Phosphorylation-independent anti-tau antibodies BR133, RD3, anti-4R and BR134, as well as phosphorylation-dependent anti-tau antibodies AT8 and AT100, were used. Two major tau of bands of 64 and 68 kDa were labelled by all antibodies, except RD3, indicating the presence of hyperphosphorylated 4R, but not 3R, tau.

**Figure 2.**
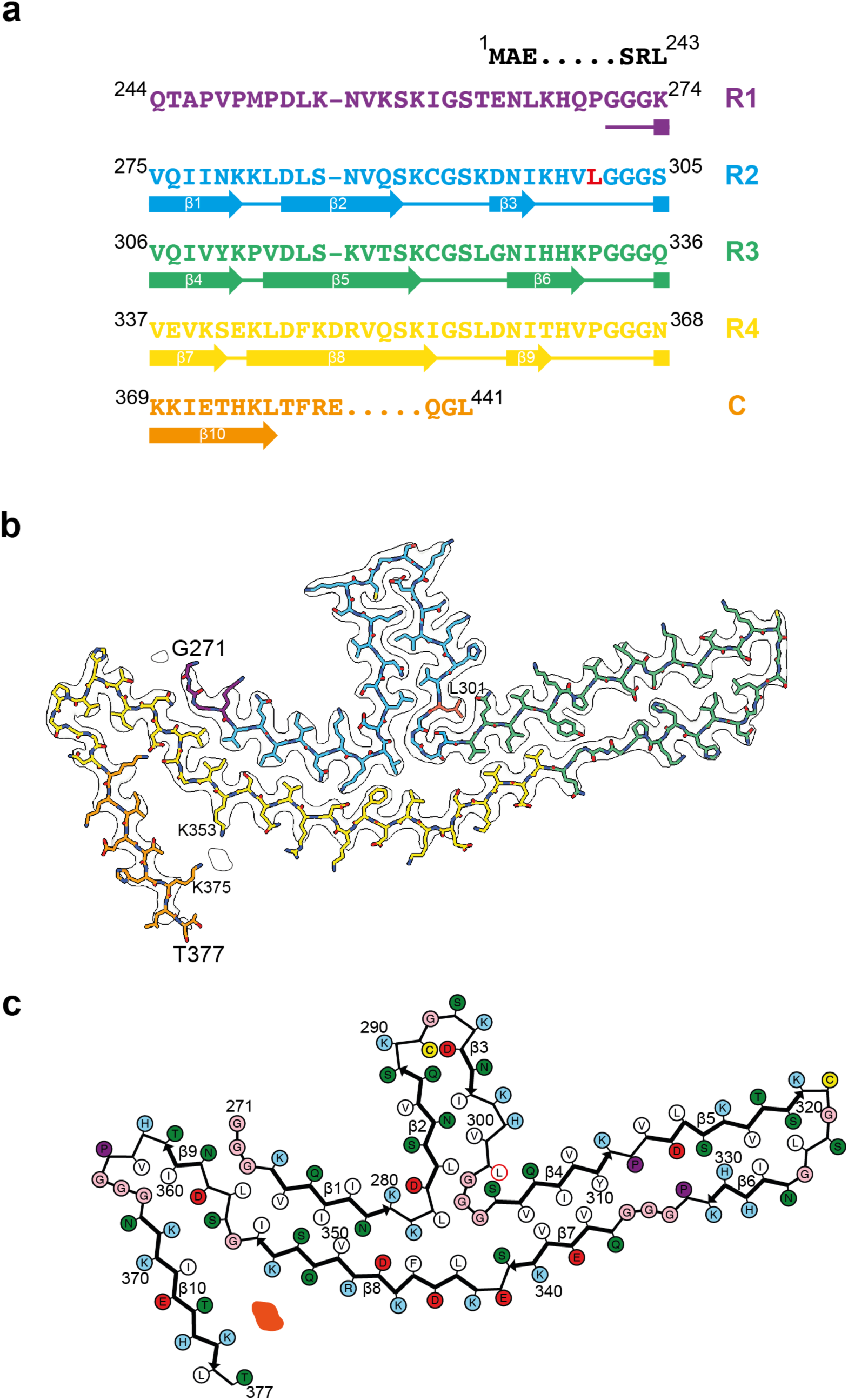
P301L mutation in *MAPT*: Cryo-EM structure of tau filaments. a, Sequence of repeats R1-R4 of tau (residues 244-368). The core structure of tau filaments extends from G271-T377 and comprises 10 β-strands (Ω1-Ω10, shown as thick arrows; loops are shown as thin lines). Residue L301 is highlighted in red. b, Sharpened cryo-EM map of tau filaments from the parietal cortex of case 2, with the atomic model overlaid. Residues in R1-R4 and the sequence after R4 are coloured purple, blue, green, gold and orange, respectively. Residue L301 is labelled, together with the N-terminal G271 and C-terminal T377 of the ordered core, as well as K353 and K375, which coordinate a non-proteinaceous density. c, Schematic of the P301L tau filament fold. Negatively charged residues are shown in red, positively charged residues in blue, polar residues in green, non-polar residues in white, sulfur-containing residues in yellow, prolines in purple and glycines in pink. Thick arrows indicate Ω-strands (Ω1-10). An internal additional density is shown in orange. Residue L301 is outlined in red.

Cryo-EM showed that the tau protofilament spans residues G271-T377 and adopts a double-layered, three-lobed fold (Figure 2). The cryo-EM density at residue 301 is consistent with a leucine side chain. Unlike the known structures of wild-type 4R tau filaments, which are folded into either three- or four-layered structures [4,34], the three-lobed fold of P301L tau consists of a stem of two-layered cross-β structure and two hairpin-like arms that are connected at a common junction. The short arm is made of residues 282-303, whereas the long arm comprises residues 304-342. The stem is composed of two segments made of residues 271-281 and 343-362, which are packed against each other. The C-terminal segment, which consists of residues 363-377 of tau, folds back onto the stem, from which it is separated by a non-proteinaceous density. The latter probably corresponds to a negatively charged cofactor of unknown identity, which is co-ordinated by the positively charged side chains of K353 and K375. It is reminiscent of the non-proteinaceous densities in the *ex vivo* structures of other human brain tau filaments [2]. At the three-lobed junction, there is a solvent-filled cavity surrounded by tau residues L282, G304, S341 and L344.

With the exception of the short arm, the three-lobed fold of P301L tau resembles the Pick fold made of 3R tau [3,35], with the conformation of the long hairpin-like arm (residues 306-341) being nearly identical to that of the same region in the Pick tau fold (Extended Data Figure 5). The similarity of the three-lobed fold to the Pick fold also extends to the stem region, where residues 347-358 adopt similar backbone conformations and form analogous interfaces with the opposite segments, which are made of residues 275-281 of the former and residues 261-267 of the latter. Residues N279 and K281 in the P301L tau fold occupy equivalent positions to residues N265 and K267 in the Pick tau fold (Extended Data Figure 5a,b).

As no structure resembling that of the short arm is present in the known filament structures of wild-type tau, its formation is probably the result of the P301L mutation stabilising the local conformation at the R2-R3 junction. Mutation of this proline results in the formation of an additional hydrogen bond between the main chains of adjacent tau molecules. The side chain of L301 is buried between the side chains of S305 and Q307 on the long arm, the polar groups of which are engaged in hydrogen bonding interactions (Figure 2b).

### Structures of tau filaments from one case of a family with mutation P301T in *MAPT*

We also determined the structures of tau filaments from the frontal cortex of a previously described individual [9,14] from a Spanish family with mutation P301T in *MAPT* (Figures 3-5). The clinicopathological diagnosis was GGT Type III. Two types of tau filaments, each made of a single protofilament, were present (Figure 3a). By immunoblotting of the sarkosyl-insoluble fraction with BR133, RD3, anti-4R, BR134, AT8 and AT100, strong tau bands of 64 and 68 kDa were observed with all antibodies except RD3, consistent with the presence of hyperphosphorylated 4R, but not 3R, tau (Figure 3b). Assembled tau was truncated at the N-terminus, as judged by the presence of additional lower molecular weight tau bands with anti-4R, BR134, AT8 and AT100, but not with BR133. By immunohistochemistry, abundant neuronal and glial 4R tau inclusions were present, with numerous globular glial tau inclusions in astrocytes and oligodendrocytes [14].

**Figure 3.**
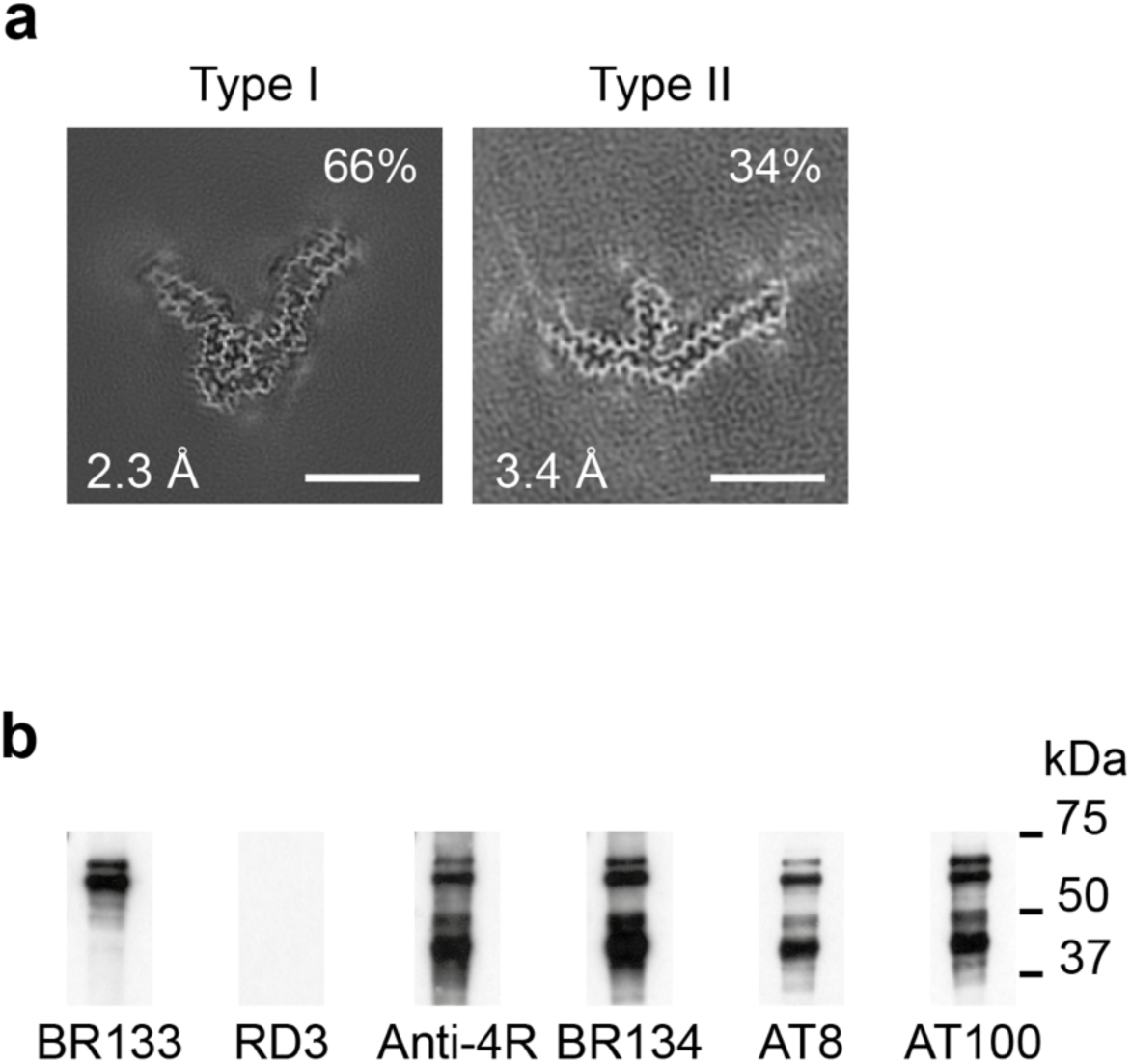
P301T mutation in *MAPT*: Cryo-EM cross-sections of tau filaments and immunoblotting. a, Cross-sections through the cryo-EM reconstructions, perpendicular to the helical axis and with a projected thickness of approximately one rung, are shown for the frontal cortex. Resolutions (in Å) and percentages of filament types are indicated at the bottom left and top right, respectively. Type I tau filaments comprised 64% and type II 36% of tau filaments Scale bars, 10 nm. b, Immunoblotting of sarkosyl-insoluble tau from the frontal cortex. Phosphorylation-independent anti-tau antibodies BR133, RD3, anti-4R and BR134, as well as phosphorylation-dependent antibodies AT8 and AT100 were used. Two major tau bands of 64 and 68 kDa were labelled by all antibodies, except RD3, indicating the presence of hyperphosphorylated 4R, but not 3R, tau.

By cryo-EM, type I filaments comprised approximately 64% and type II filaments around 36% of tau filaments (Figure 3a). Even though both types of filaments are made of single protofilaments, their ordered cores have different folds. In type II P301T tau filaments, the core spans residues N269-P364 and adopts a three-lobed fold resembling that of P301L tau filaments (Figure 4; Extended Data Figure 5a). This fold also consists of a stem and two hairpin-like arms, but it is shorter than the P301L fold at the C-terminus. Its long arm (residues 304-342) is practically identical to that of P301L tau filaments (Extended Data Figure 5b), but the short arm and the stem differ by their relative orientations and the packing of interior residues. In the stem, the N-terminal segment (residues 271-281) is shifted by two residues relative to the C-terminal segment (residues 343-364), forming an interface resembling that between these segments in the P301S tau filament fold of PS19 transgenic mice [32]. In the short arm, the opposite sides of the hairpin-like structure of type II P301T filaments have conformations resembling those of P301L tau filaments, but they are shifted relative to each other by two residues. These differences correlate with a local conformational difference around the mutation site at residue 301, the cryo-EM density of which is consistent with that of threonine. The side chain of T301 fits inside the turn formed by the glycine triplet (residues 302-304) and hydrogen bonds with S305 (Figure 4b). Compared to L301, T301 is located on the other side of the side chain of S305.

**Figure 4.**
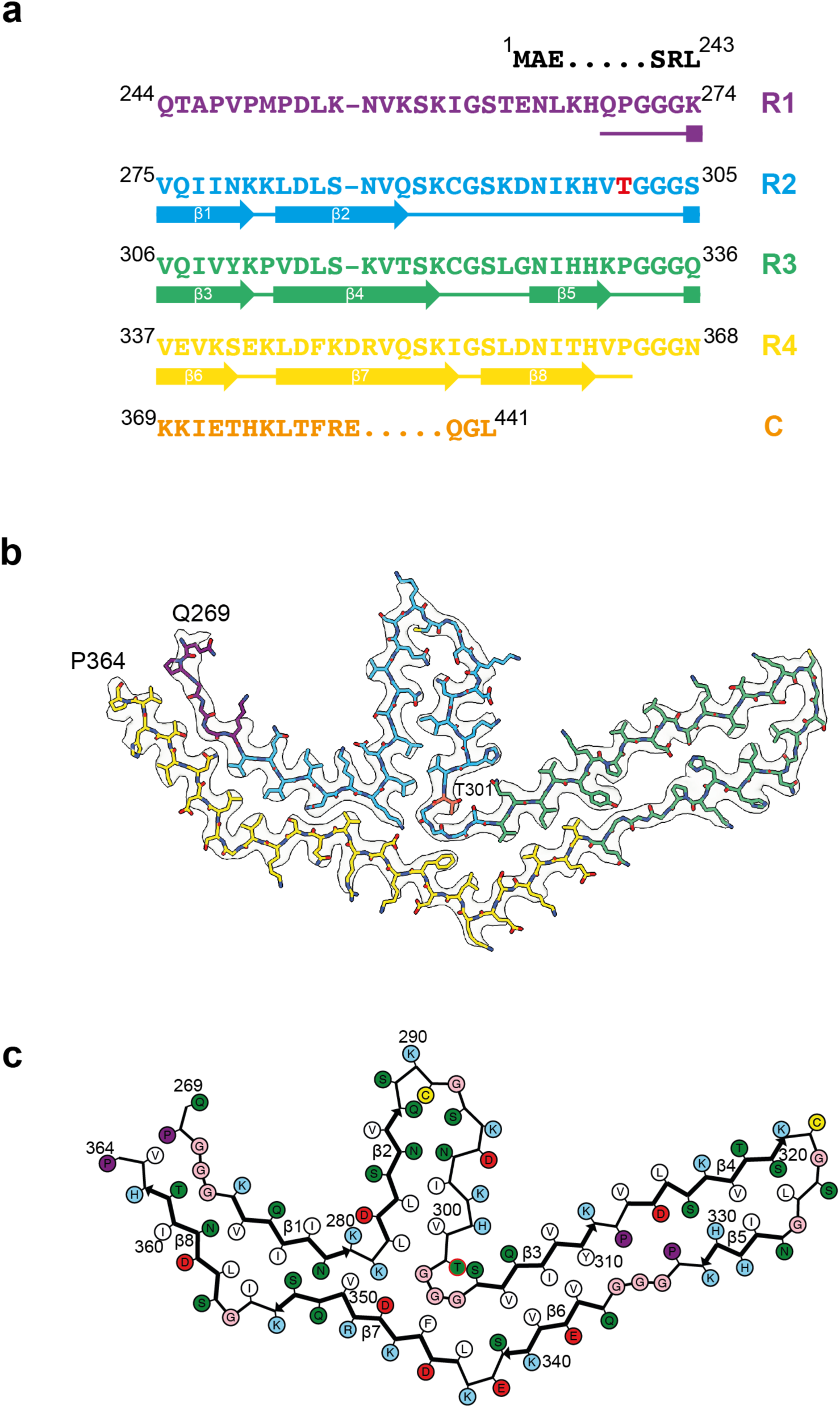
P301T mutation in *MAPT*: Cryo-EM structure of type II tau filaments. a, Sequence of repeats R1-R4 of tau (residues 244-368). The core structure of tau filaments extends from N269-P364 and comprises 8 β-strands (β1-β8, shown as thick arrows; loops are shown as thin lines). Residue T301 is highlighted in red. b, Sharpened cryo-EM map of type II tau filaments from the frontal cortex, with the atomic model overlaid. Residues in R1-R4 and the sequence after R4 are coloured purple, blue, green, gold and orange, respectively. Residue T301 is labelled and so are N-terminal residue N269 and C-terminal residue P364 of the ordered core. c, Schematic of the type II P301T tau filament fold. Negatively charged residues are shown in red, positively charged residues in blue, polar residues in green, non-polar residues in white, sulfur-containing residues in yellow, prolines in purple and glycines in pink. Thick arrows indicate β-strands 1-8. The additional density is shown in orange. Residue T301 is outlined in red.

In type I P301T filaments, the core spans residues G273-R379 of tau and adopts a new fold, in the shape of the letter V (Figure 5). It has a four-layered body and two-layered wings. The N-terminal half of the core sequence (residues 273-324) forms the inner side of the V-shaped fold with a sharp bend at the R2-R3 junction, whereas the C-terminal half (residues 325-379) curves around this bend on the outer side.

**Figure 5.**
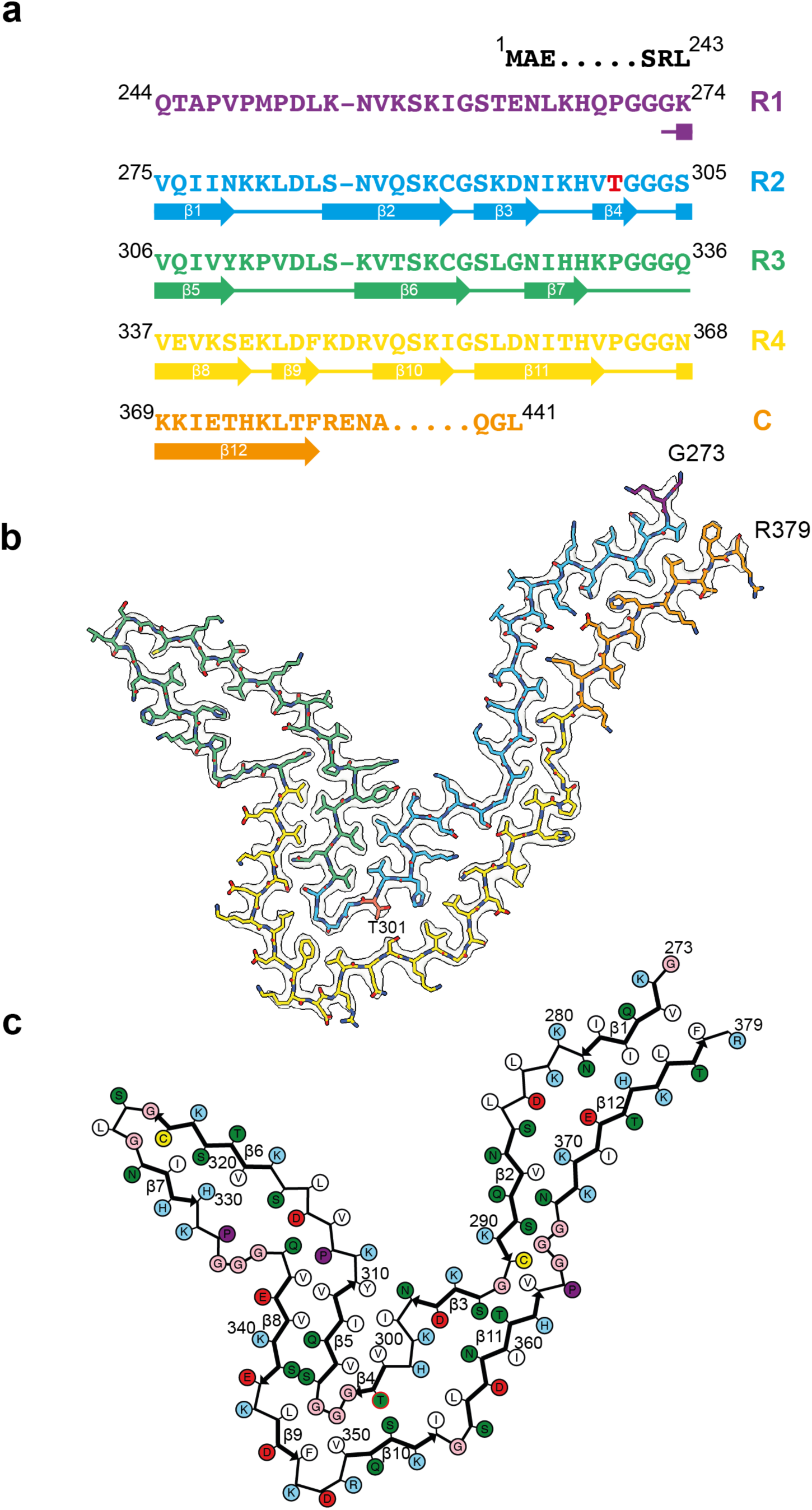
P301T mutation in *MAPT*: Cryo-EM structure of type I tau filaments. a, Sequence of repeats R1-R4 of tau (residues 244-368). The core structure of type I tau filaments extends from G273-R379 and comprises 12 β-strands (β1-β12, shown as thick arrows; loops are shown as thin lines). Residue T301 is highlighted in red. b, Sharpened cryo-EM map of type I tau filaments from the frontal cortex, with the atomic model overlaid. Residues in R1-R4 and the sequence after R4 are coloured purple, blue, green, gold and orange, respectively. c, Schematic of the type I P301T tau filament fold. Negatively charged residues are shown in red, positively charged residues in blue, polar residues in green, non-polar residues in white, sulfur-containing residues in yellow, prolines in purple and glycines in pink. Thick arrows indicate β-strands 1-12.

The conformation of the R2-R3 junction in type I P301T filaments is different from that of type II P301T filaments, but it is almost identical to the conformations of the R2-R3 junction in filaments from mouse lines expressing human tau with mutations P301S or P301L (Extended Data Figure 6) [32,33]. A similar conformation of this junction was also observed in the GPT fold of wild-type 4R tau [4]. Missense mutations in P301 stabilise this conformation by creating an additional hydrogen bond to the main chain of an adjacent molecule. The side chain of T301 is on the outside surface of this common substructure and makes a solvent-mediated interaction with residue S352 of R4.

The type I P301T fold also displays local similarities with the corticobasal degeneration (CBD) and AGD tau folds by sharing with them the substructures 312-333 in R3 and 337-357 in R4. In all three folds, residues 312-333 form a compact semi-detached unit, whereas residues 337-357 wrap around the R2-R3 junction. However, in the type I P301T filaments, the substructure from R4 makes interactions with the R2-R3 junction that have an inside-out conformation compared to the CBD and AGD folds. In addition, in the type I P301T and in the CBD folds, there are similar interfaces between the N-terminal (residues 275-281) and C-terminal (residues 372-378) segments, which are made of the same residues, but are shifted relative to each other.

## Discussion

FTDP-17 is a heterogeneous clinicopathological entity, with the long arm of chromosome 17 harbouring both the progranulin gene (*GRN*) and *MAPT*. Heterozygous loss-of-function mutations in *GRN* give rise to FTDP-17 with abundant TDP-43 inclusions [37–40], whereas heterozygous gain-of-toxic-function mutations in *MAPT* give rise to FTDP-17 with abundant tau inclusions [6,7,41]. These different molecular constituents of the filamentous inclusions suggest that *GRN* and *MAPT* mutations lead to different diseases. We have previously shown that cases of FTDP-17 caused by *MAPT* mutations can be subdivided and that the different filament folds of mutant tau are identical to those of sporadic tauopathies [3–5]. The structures presented here, which are the first tau folds of FTDP-17 that are different from those of sporadic tauopathies, provide additional insights into this heterogeneity.

The atomic structures of tau filaments from the brains of five individuals with a P301L tau mutation belonging to three unrelated families and the structures of tau filaments from the brain of an individual with a P301T mutation are distinct from any of the structures described previously for individuals with *MAPT* mutations, including V337M, R406W, ΔK281 and intron 10 mutations +3 and +16 [3–5]. Whereas mutations V337M and R406W lead to the same tau filaments as in AD [5], ΔK281 gives rise to tau filaments with the Pick fold [3], and intron 10 mutations +3 and +16 lead to filaments with the AGD fold [4]. Mutations P301L and P301T in tau give rise to new folds that have not been described previously for any of the sporadic or familial tauopathies.

So far, specific tau folds have defined different diseases, even though the same fold can be found in multiple conditions, such as the chronic traumatic encephalopathy (CTE) fold in CTE, subacute sclerosing panencephalitis (SSPE), amyotrophic lateral sclerosis/parkinsonism-dementia complex (ALS/PDC) and vacuolar tauopathy (VT) [2,42]. If the same is true of FTDP-17, it follows that the finding that tau filament structures in tauopathies caused by mutations P301L and P301T are novel, implies that these cases are separate familial tauopathies. Moreover, tau filaments from an individual with mutation P301T tau, which gives rise to GGT type III [14], have folds that are distinct from those of sporadic GGT types I/II [4]. We previously reported that the tau filaments from a case of sporadic GGT type III are straight, precluding the determination of their structures [4]. GGT type III may thus be a different disease from types I/II and filament structures may differ between sporadic and inherited forms of disease. Cases of P301L tau with a clinicopathological diagnosis of GGT have been described [11–13], but their tau filament structures remain to be determined.

Like the tau filaments from progressive supranuclear palsy (PSP), all the tau filaments from individuals with P301L and P301T tau were made of single protofilaments. While we observed the same tau fold in filaments extracted from the cerebral cortex of five individuals with mutation P301L tau, we had only access to brain material from a single individual with mutation P301T, for whom we observed two different protofilament folds. Intriguingly, the type II P301T fold resembled the P301L tau fold. It will be interesting to see if additional cases with mutation P301T in *MAPT* have the same folds as those shown here.

Surprisingly, the three-lobed folds of 4R tau with the P301L or P301T folds bear similarity to the Pick fold of 3R tau [35]. In human cases with the P301L tau mutation, characteristic mini pick-like bodies have been described by light microscopy, especially in the dentate gyrus [11,43,44]. However, unlike Pick bodies [45], the 4R tau inclusions from an individual with mutation P301L tau were Gallyas-Braak silver-positive.

Filaments of tau from the brains of individuals with a P301 mutation are made only of mutant tau, unlike filaments from individuals with mutations V337M and R406W, which adopt the Alzheimer tau fold and can consist of a mixture of wild-type and mutant proteins [5]. Wild-type human tau may be unable to form the P301L or P301T folds, even in the presence of seeds made of either P301L or P301T tau. The observation that the P301L tau folds are different from the folds of wild-type tau may explain why PET ligand [^18^F]-flortaucipir, which binds with high affinity to the Alzheimer tau fold, showed only little binding in individuals with mutation P301L in *MAPT* [46–49].

Different conformations of the R2-R3 junctions in the P301L and P301T tau folds of filaments extracted from human brains, together with work on filaments extracted from the brains of mice transgenic for human P301S or P301L tau [32,33], underscore the polymorphic nature of filaments assembled from tau with a P301 mutation. The structures of P301L tau filaments from human brains are different from those of P301L tau filaments from transgenic mouse line rTg4510 [34]. The latter also differ from the structures of tau filaments from mouse lines expressing human mutant P301S tau [32].

The lengths of tau filament cores from transgenic mouse lines rTg4510, Tg2541 and PS19 are shorter than those of tau filaments extracted from human brains [2,32,33]. Unlike tau filaments from human brains, mouse brain tau filament cores do not contain the whole of R4 or the sequence after R4. Of note, the P301L tau filaments from mouse line rTg4510 and the P301S tau filaments from line Tg2541 share a large substructure outside the mutation site that has not been observed in the tau filaments extracted from human brains. The structures of tau filaments from humans with P301S tau are not known. It remains to be determined if transgenic mouse lines can be produced that form the same tau filament folds as those found in human brains.

Mutation of P301 in tau has led to the development of transgenic mouse models that develop hyperphosphorylation of tau, filament formation and neurodegeneration [25–29]. However, the observed polymorphisms in tau filaments with P301 mutations suggest that such models should be used with caution in the study of tau pathologies of either AD or sporadic 4R tauopathies, such as PSP.

## Methods

### Cases with *MAPT* mutation P301L

We used parietal cortex from two individuals (cases 1 and 2) belonging to two separate US families with mutation P301L in *MAPT*. Case 1 was a female who died aged 62 after a 10-year history of personality changes and cognitive impairment. Her mother died with a dementing illness and her sister has FTD. Case 2 was a female who died aged 55 after an 11-year history of FTD. No family history of dementia was reported. We also used also temporal cortex from three individuals (cases 3-5) belonging to a large pedigree (family 1) from the Netherlands with mutation P301L in *MAPT* [7,50]. A variety of symptoms developed, consistent with a diagnosis of behavioural-variant FTD. Case 3 was a male who died aged 55 after a 2-year history of FTD. His mother, brother and sister had suffered from FTD. Case 4 was a male who died aged 57 after a 4-year history of FTD. His mother and two of his brothers had FTD. Case 5 was a male who died aged 64 after an 11-year history of FTD. His mother, her brother and her sister suffered from FTD.

### Case with *MAPT* mutation P301T

We used frontal cortex from a previously described male (case III.3.1. in [9]; case 1 in [14]) from a Spanish family with mutation P301T in *MAPT.* This individual had a C to A nucleotide substitution in the first position of codon 301 (CCG to ACG, on one allele). At age 45, he started to develop a pyramidal syndrome with features of primary lateral sclerosis, severe supranuclear ophthalmoplegia, parkinsonism and dementia. He died aged 49 with a clinical diagnosis of progressive supranuclear palsy with primary lateral sclerosis. The clinicopathological diagnosis was GGT type III. The subject’s mother died aged 70 with a clinical diagnosis of probable corticobasal degeneration and his maternal grandfather suffered from dementia and died aged 53.

### Genomic analysis

Genomic DNA was extracted from brain or from blood with informed consent. Standard amplification reactions were done with 50 ng genomic DNA, followed by DNA sequencing of exons 1 and 9-13 of *MAPT* with adjoining intronic sequences, as described [51]. The relatedness of individual cases was determined using whole exome sequencing.

### Filament extraction

Sarkosyl-insoluble material was extracted from parietal cortex of P301L tau cases 1 and 2, temporal cortex of P301L tau cases 3-5 and frontal cortex of the P301T tau case, as described [52]. Tissues were homogenised in 20 vol (w/v) buffer A (10 mM Tris-HCl, pH 7.4, 0.8 M NaCl, 10% sucrose and 1 mM EGTA), brought to 2% sarkosyl and incubated at 37° C for 30 min. The samples were centrifuged at 10,000 g for 25 min, followed by spinning of the supernatants at 100,000 g for 60 min. The pellets were resuspended in buffer A (700 μl/g tissue) and centrifuged at 5,000 g for 5 min. The supernatants were diluted 3-fold in buffer B (50 mM Tris-HCl, pH 7.5, 0.15 M NaCl, 10% sucrose and 0.2% sarkosyl), followed by a 30 min spin at 166,000 g. For cryo-EM, the pellets were resuspended in 150 μl/g buffer C (20 mM Tris-HCl, pH 7.4, 100 mM NaCl).

### Immunoblotting and histology

For immunoblotting, samples were resolved on 4-12% Bis-Tris gels (NuPage) and the primary antibodies [BR133, 1:4,000; RD3 (Sigma-Millipore, 1:4,000); anti-4R (Cosmo-Bio, 1:2,000); BR134, 1:4,000; AT8 (Thermo Fisher, 1:11,000); AT100 (Thermo Fisher, 1:500)] were diluted in PBS plus 0.1% Tween-20 and 5% non-fat dry milk. Histology and immunohistochemistry were carried out as described [50]. Some sections (8 μm) were counterstained with haematoxylin-eosin. The primary antibodies were: RD3 (1:3,000); anti-4R (1:400); AT8 (1:1,000).

### Electron cryo-microscopy

Cryo-EM grids (Quantifoil 1.2/1.3, 300 mesh) were glow-discharged for 1 min using an Edwards (S150B) sputter coater. Three μl of the sarkosyl-insoluble fractions was applied to the glow-discharged grids, followed by blotting with filter paper and plunge freezing into liquid ethane using a Vitrobot Mark IV (Thermo Fisher Scientific) at 4° C and 100% Krios G2 or G4 microscope (Thermo Fisher Scientific) operated at 300 kV. For P301L tau case 5, movies were acquired on a Gatan K2 summit detector using a pixel size of 1.1 Å. For P301L tau cases 3 and 4, images were acquired on a Gatan K3 detector with a pixel size of 0.93 Å. For both detectors, a quantum energy filter with a slit width of 20 eV was used to remove inelastically scattered electrons. For P301L tau cases 1 and 2 and for the P301T tau case, movies were acquired on a Falcon-4 detector using a pixel size of 0.824 Å. Images were recorded in electron event representation format [53]. See Extended Data Table 1 and Extended Data Figure 7 for further detail.

### Data processing

Datasets were processed in RELION using standard helical reconstruction [54,55]. Movie frames were gain-corrected, aligned and dose-weighted using RELION’s own motion correction programme [56]. Contrast transfer function (CTF) was estimated using CTFFIND4.1 [57]. Filaments were picked manually. Initial models were generated *de novo* from 2D class average images using relion_helix_inimodel2d [58]. Three-dimensional refinements were performed in RELION-4.0 and the helical twist and rise refined using local searches. Bayesian polishing and CTF refinement were sharpened using post-processing procedures in RELION-4.0 [59] and resolution estimates were calculated based on Fourier shell correlation (FSC) between two independently refined half-maps at 0.143 (Extended Data Figure 6) [60]. We used relion_helix_toolbox to impose helical symmetry on the post-processing maps.

### Model building and refinement

Atomic models of the P301L and P301T (type I and type II) filaments were built manually using Coot [61]. Refinements were performed using ISOLDE [62], *Servalcat* [63] and REFMAC5 [64,65]. Models were validated with MolProbity [66]. Figures were prepared with ChimeraX [67] and PyMOL [68]. When multiple maps of the same filament type were resolved, atomic modelling and database submission were only performed for the map with the highest resolution.

## Acknowledgements

We thank the patients’ families for donating brain tissues, T. Darling, I. Clayson and J. Grimmett for high-performance computing and the EM facility of the Medical Research Council (MRC) Laboratory of Molecular Biology for help with cryo-EM data acquisition. We are grateful to R. Richardson, N. Maynard, M. Jacobsen and B. Glazier for help with histology and immunohistochemistry. This work was supported by the MRC, as part of U.K. Research and Innovation (UKRI) (MC_UP_A025_1013 to S.H.W.S. and MC_1051284291 to M.G.). It was also funded by the National Natural Science Foundation of China (82371415, to Y.S.) and the Key R&D Program of Zhejiang (2024SSYS0018, to Y.S.). This work was also supported by the U.S. National Institutes of Health (P30-AG010133, R01-AG080001 and RF1-AG071177, to R.V. and B.G.) and the Department of Pathology and Laboratory Medicine, Indiana University School of Medicine (to R.V. and B.G.).

## Author contributions

E.E., H.S., I.F., J.C.V. and B.G. identified patients and performed neuropathology; H.J.G., R.V. and J.R.M. performed genetic analysis; M.S. performed immunoblotting; M.S. and Y.S. collected cryo-EM data; M.S., Y.S. and A.G.M. analysed cryo-EM data.; S.H.W.S. and M.G. supervised the project. All authors contributed to the writing of the manuscript.

## Competing interests

The authors have no competing interests.

## Data availability

Cryo-EM maps have been deposited in the Electron Microscopy Data Bank (EMDB) with the accession numbers EMD-51319 for P301L tau filaments from case 2, EMD-51320 for type I P301T tau filaments and EMD-51325 for type II P301T tau filaments. The corresponding refined atomic models have been deposited in the Protein Data Bank (PDB) under accession numbers 9GG0 for P301L tau filaments from case 2, 9GG1 for type I P301T tau filaments and 9GG6 for type II P301T tau filaments.

## Extended Data Figure Legends

**Extended Data Figure 1.**
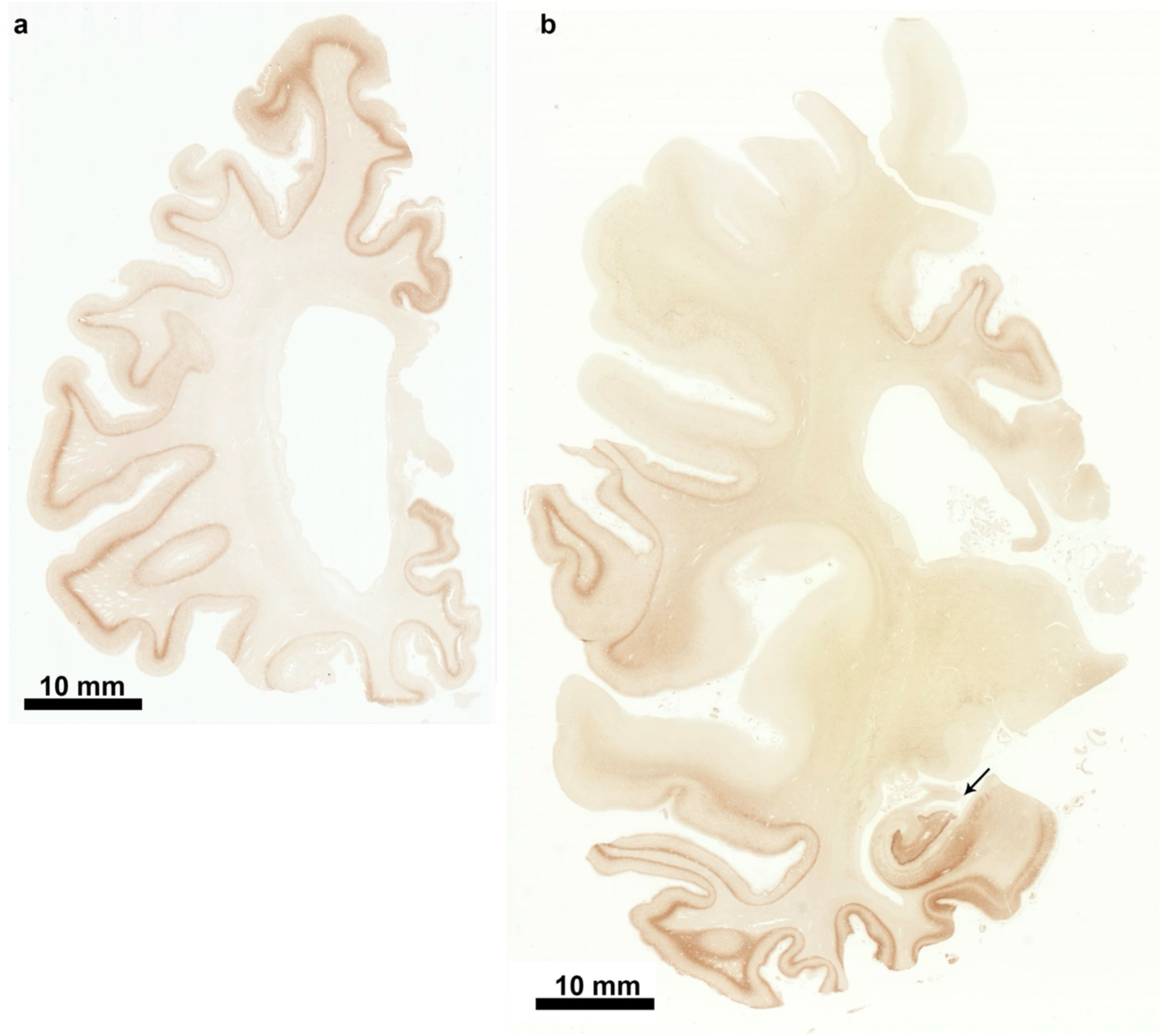
P301L mutation in *MAPT*: Tau inclusions in coronal brain sections from case 1. a, Coronal section of frontal lobe stained with phosphorylation-dependent anti-tau antibody AT8. Note that all gyri are strongly stained. Scale bar, 10 mm. b, Coronal section of parietal and temporal lobes stained with phosphorylation-dependent anti-tau antibody AT8. Note that not all gyri are labelled in the parietal lobe (upper part of the section). In the temporal lobe, inferior gyri and the hippocampus (arrowed) are strongly stained (lower part of the section). Scale bar, 10 mm.

**Extended Data Figure 2.**
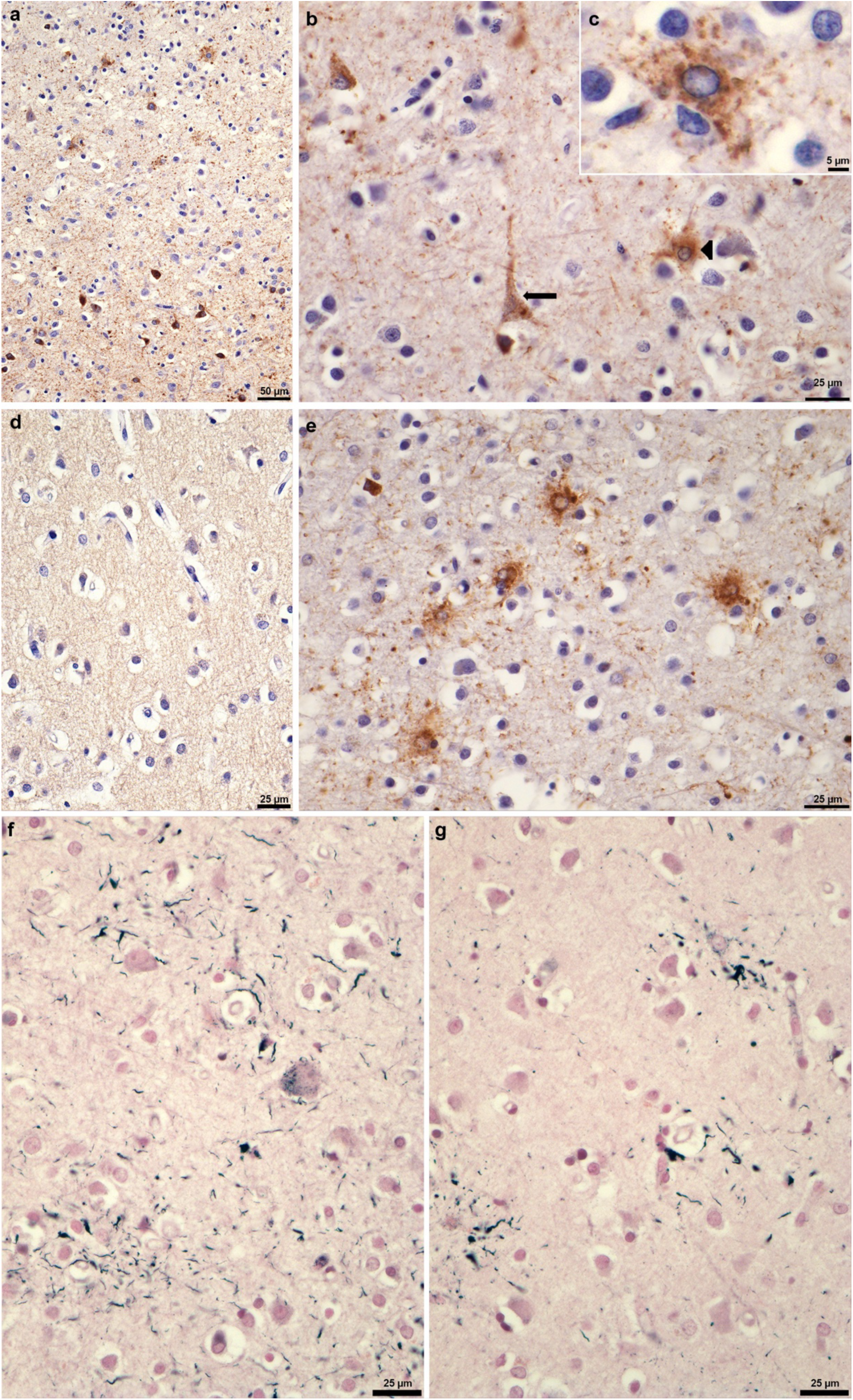
P301L mutation in *MAPT*: Tau inclusions in parietal cortex (postcentral gyrus) from case 1. The following anti-tau antibodies were used: AT8 (a,b,c); RD3 (d); RD4 (e). Gallyas-Braak silver staining was used in (f,g). a, AT8-positive nerve cells (lower part of the image) and astrocytes (upper part of the image). Scale bar, 50 μm. b, AT8-positive nerve cell (arrow) and AT8-positive astrocyte (arrowhead). Cell bodies and processes are labelled. Scale bar, 25 μm. c, An astrocyte with AT8-positive cytoplasm and processes and AT8-negative nucleus. Scale bar, 5 μm. d, No labelling of neurons or astrocytes by RD3. Scale bar, 25 μm. e, Astrocytes are labelled by RD4 (this case had more tau-immunoreactive astrocytes than neurons). Scale bar, 25 μm. f, Nerve cells and their processes are Gallyas-Braak silver-positive. Scale bar, 25 μm. g, Astrocytic processes are Gallyas-Braak silver-positive. Scale bar, 25 μm.

**Extended Data Figure 3.**
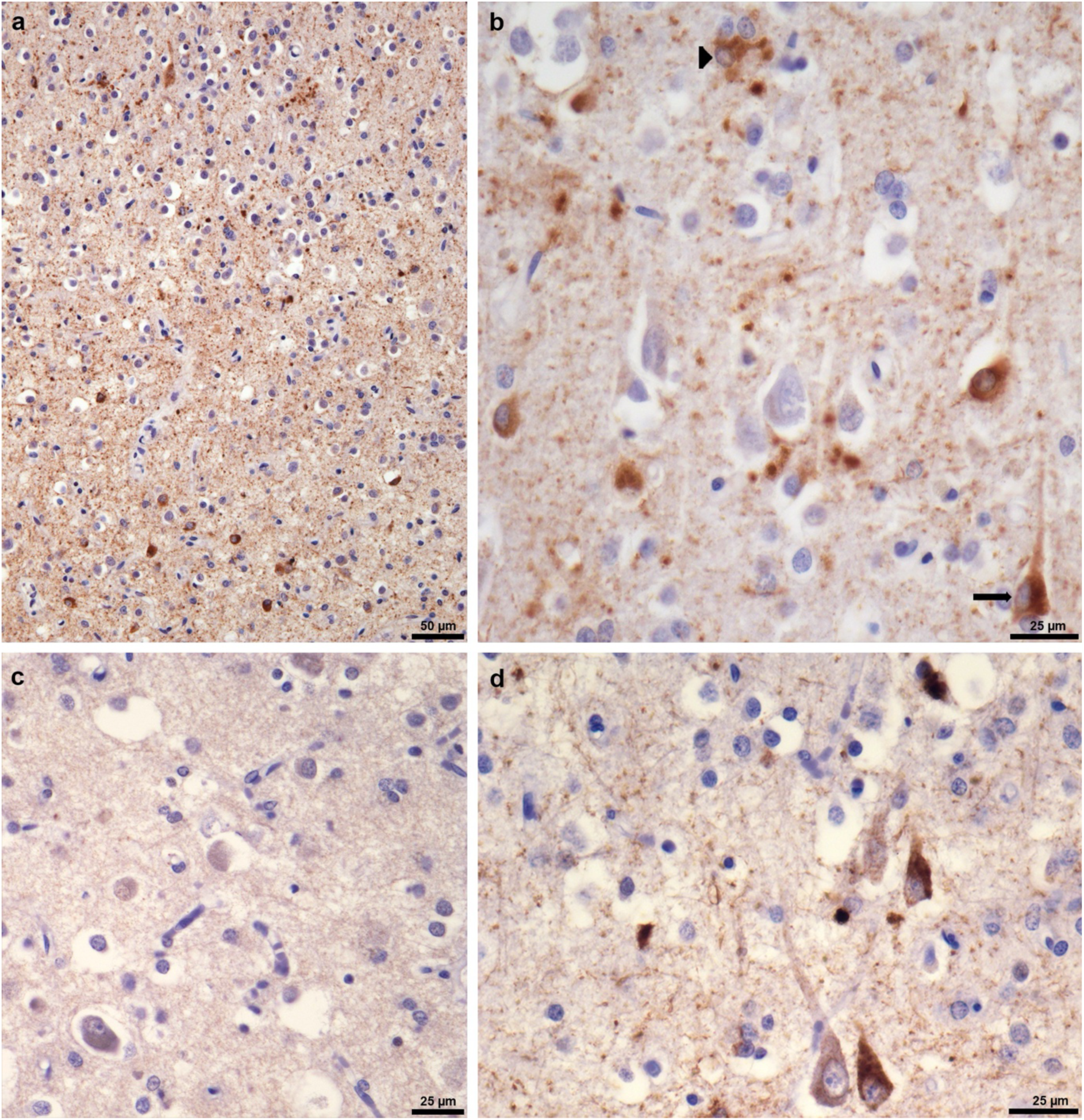
P301L mutation in *MAPT*: Tau inclusions in parietal cortex (postcentral gyrus) from case 2. The following anti-tau antibodies were used: AT8 (a,b); RD3 (c); RD4 (d). a, AT8-positive nerve cells (lower part of the image) and astrocytes (upper part of the image). Scale bar, 50 μm. b, AT8-positive nerve cell (arrow) and AT8-positive astrocyte (arrowhead). Some neuropil elements are also AT8-immunoreactive. Scale bar, 25 μm. c, No labelling of neurons or astrocytes by RD3. Scale bar, 25 μm. d, Nerve cells and neuropil threads are labelled by RD4 (this case had more tau-immunoreactive neurons than astrocytes). Scale bar, 25 μm.

**Extended Data Figure 4.**
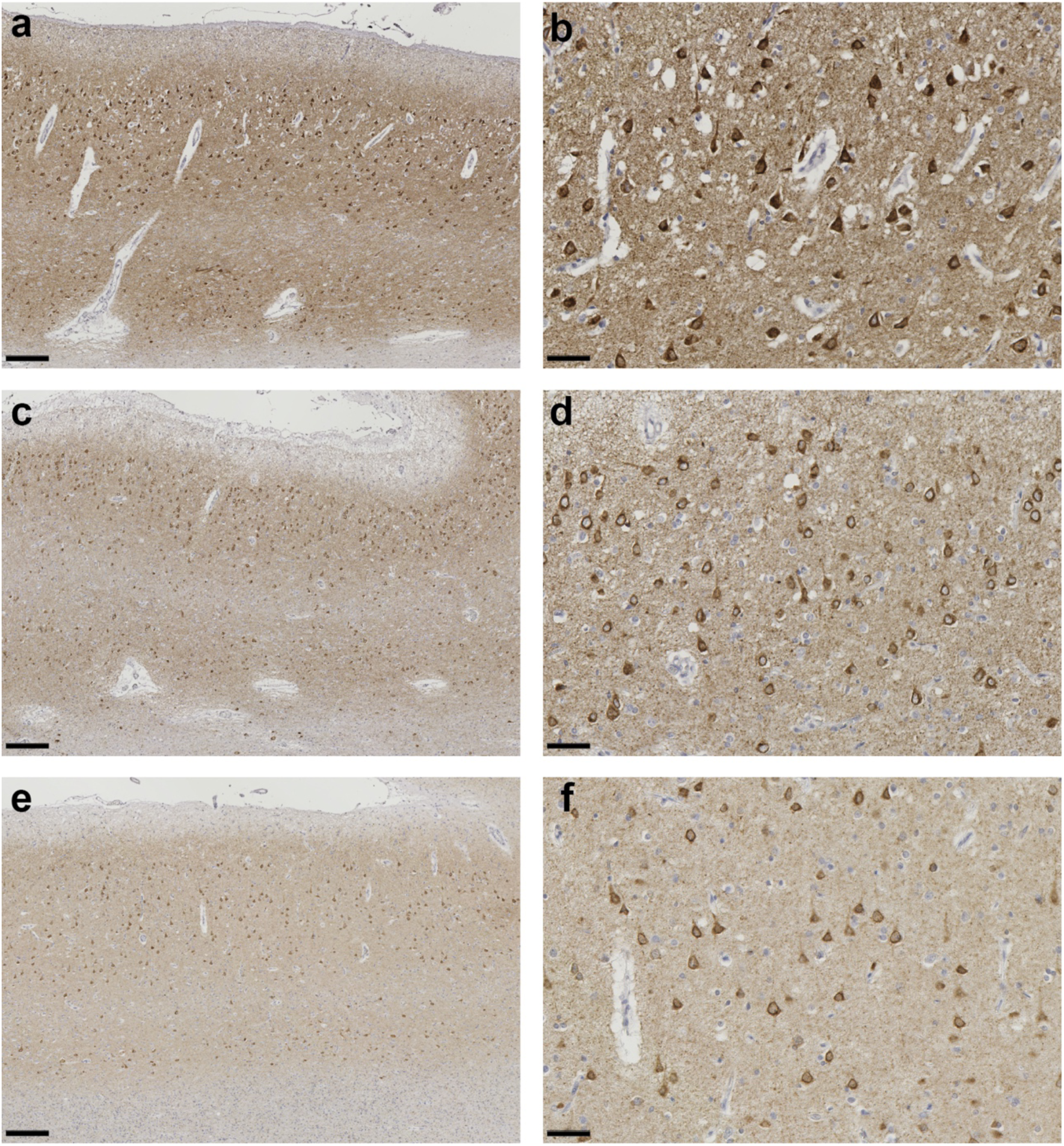
P301L mutation in *MAPT*: Tau inclusions in temporal cortex from cases 3-5. (a-f), Staining with phosphorylation-dependent anti-tau antibody AT8, showing numerous positive neuronal and glial cell inclusions in P301L tau cases 3 (a,b), 4 (c,d) and 5 (e,f). Scale bars: 200 μm (a,c,e); 50 μm (b,d,f).

**Extended Data Figure 5.**
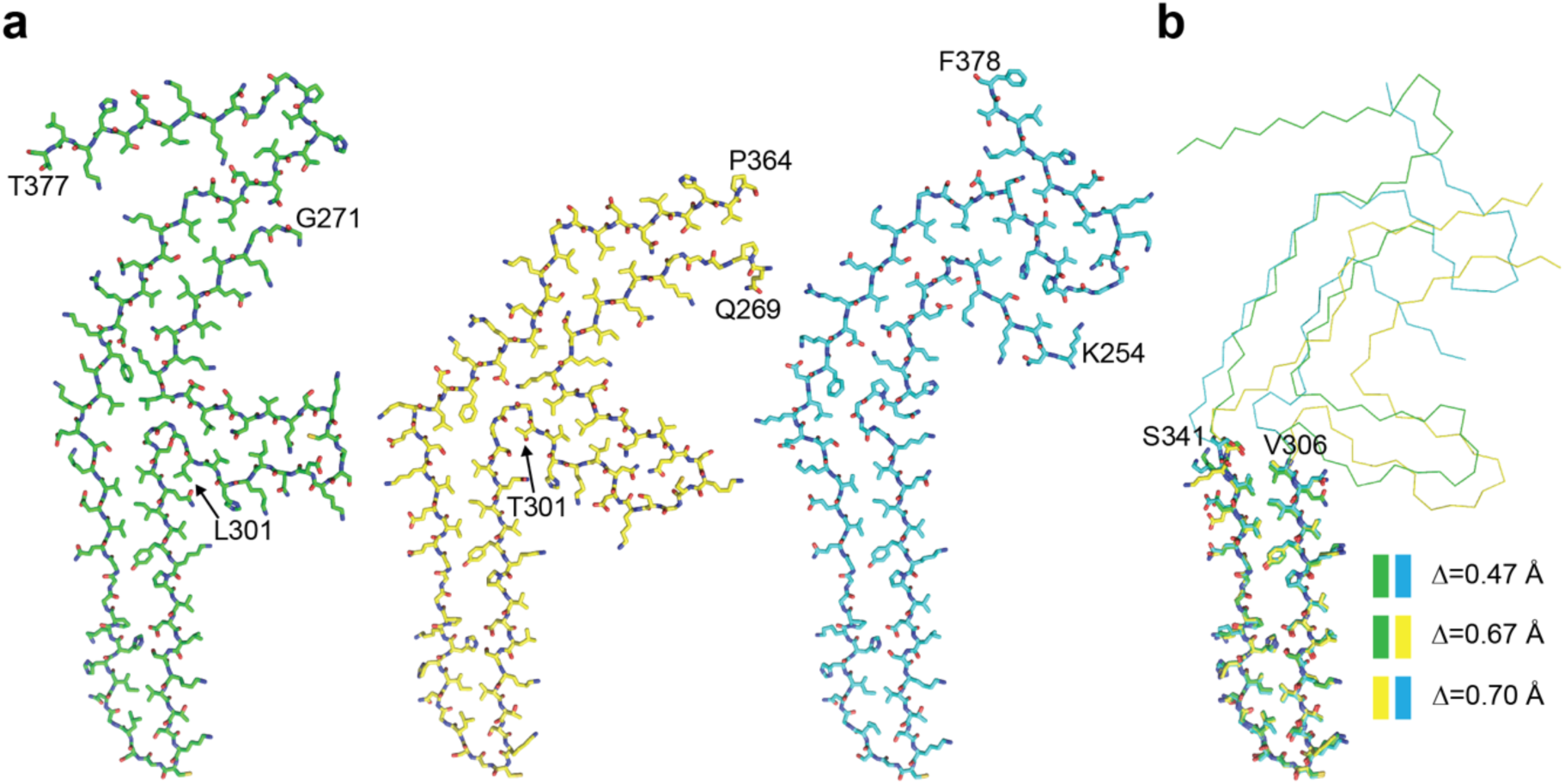
Comparison of the three-lobed folds of P301L and P301T tau filaments with the Pick tau fold. a, Side-by-side comparison of the three-lobed folds of P301L (left, green) and type II P301T (middle, yellow) tau filaments with the Pick tau fold (right, cyan) (PDB:8P34). b, Overlay of the long arm substructures (residues V306-S341) shown as sticks. Backbone root mean square deviation values Δ were calculated for each pair of substructures (colour coded as in panel a). The remainder of the three folds is shown as backbone.

**Extended Data Figure 6.**
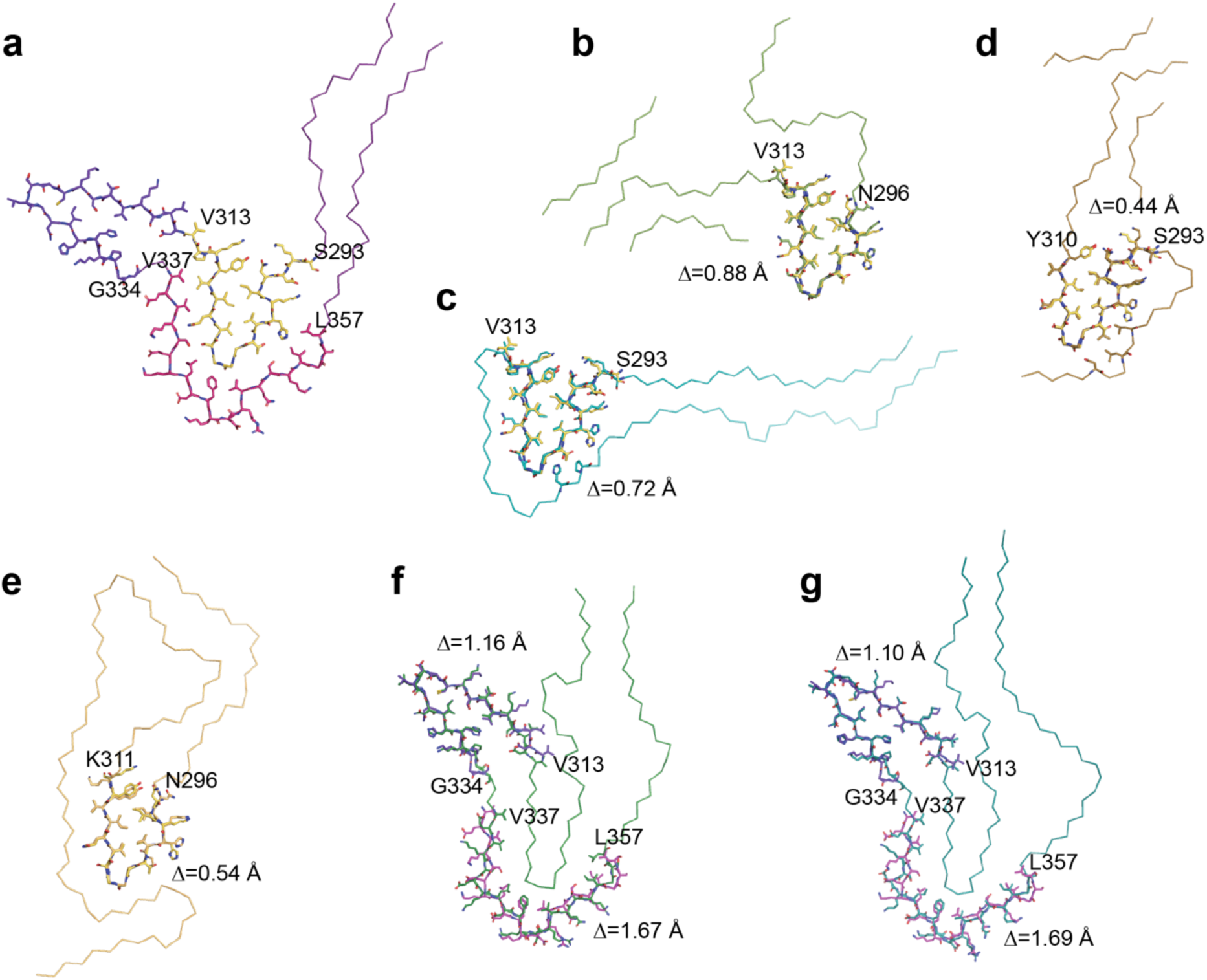
Comparison of the type I P301T tau fold with structures of tau filaments from human brains and transgenic mouse brains. a, Ribbon plot of the type I P301T fold. Shown as sticks and highlighted by different colours are substructures shared with other filaments. b-g, Overlays of the corresponding substructures are shown as sticks. Backbone root mean square deviation values Δ are shown for each overlay. b, P301S tau filament fold from mouse line Tg2541 (PDB:8Q96). c, P301S tau filament fold from mouse line PS19 (PDB:8Q92). d, P301L tau filament fold from mouse line rTg4510 (PDB:8WCP). e, Globular glial tauopathy-progressive supranuclear palsy tau (GPT) fold (PDB:7P6A). f, Corticobasal degeneration (CBD) fold (PDB:6TJO). g, Argyrophilic grain disease (AGD) fold (PDB:7P6D).

**Extended Data Figure 7.**
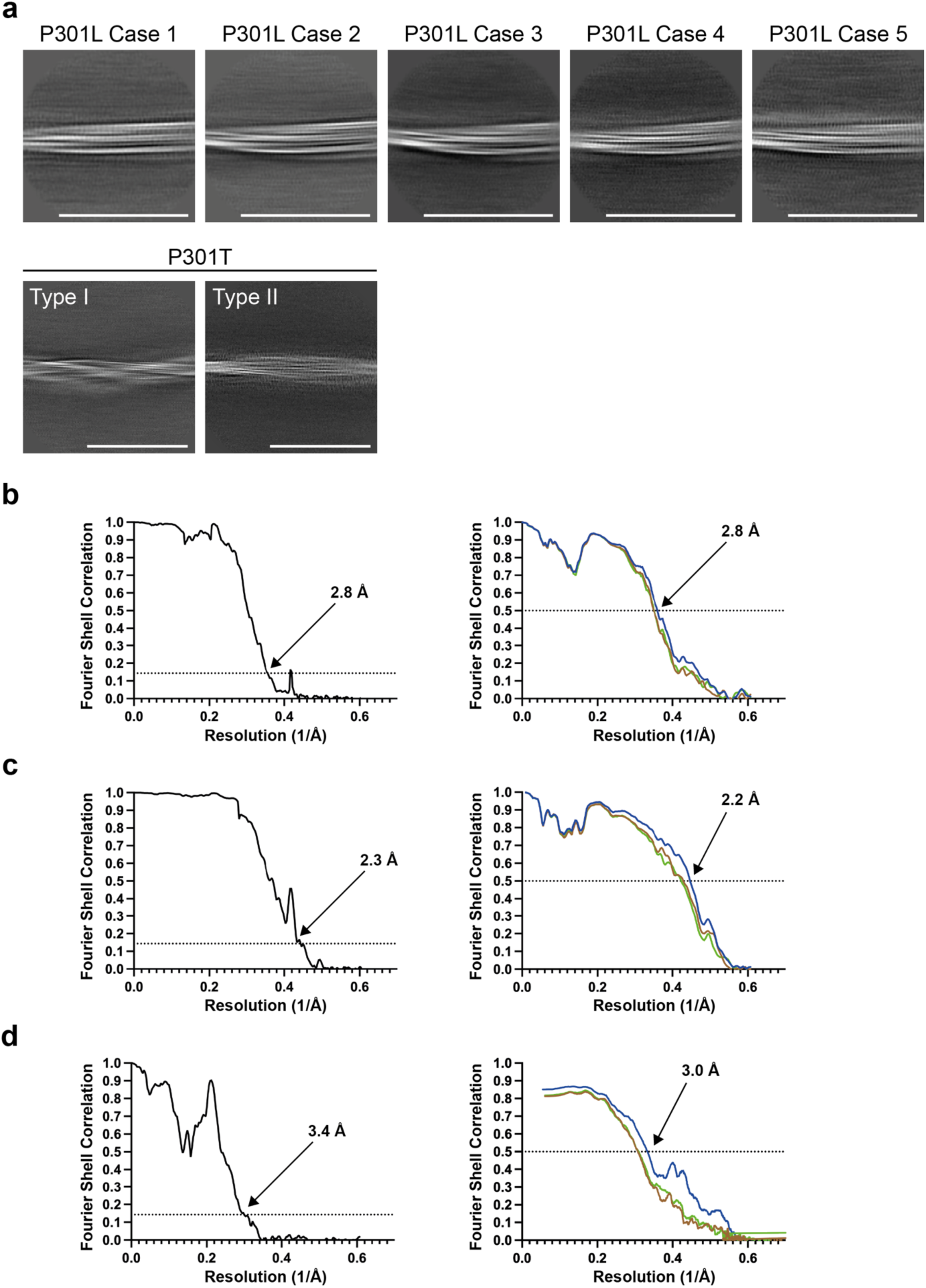
Cryo-EM 2D classification and resolution estimates. a, Representative 2D classification images of tau filaments from cases with *MAPT* mutations P301L and P301T. Scale bars, 50 nm. b-d, Solvent-corrected Fourier shell correlation (FSC) curves of cryo-EM half-maps (left panels) and model-to-map validation (right panels) for P301L tau filaments (from case 2) (b), P301T tau filaments type I (c) and P301T tau filaments type II (d). FSC curves for the final refined atomic model against the final cryo-EM map are shown in red; for the atomic model refined in the first half map in brown (model 1 versus half map 1); and for the refined atomic model in the first half map against the other half map in green (model 1 versus half map 2).

**Extended Data Table 1.**

|  | P301L<br>Case 1 | P301L<br>Case 2 | P301L<br>Case 3 | P301L<br>Case 4 | P301L<br>Case 5 | P301T<br>Case 1 |  |
| --- | --- | --- | --- | --- | --- | --- | --- |
| Data collection and processing |  |  |  |  |  |  |  |
| Magnification | 96,000 | 96,000 | 81,000 | 81,000 | 105,000 | 96,000 |  |
| Voltage (kV) | 300 | 300 | 300 | 300 | 300 | 300 |  |
| Detector | Falcon 4 | Falcon 4 | K3 | K3 | K2 | Falcon 4 |  |
| Electron dose (e−/Å²) | 30.0 | 30.0 | 32.7 | 33.3 | 42.5 | 42.0 |  |
| Defocus range (µm) | 1.8-2.4 | 1.7-2.4 | 1.6-2.4 | 1.6-2.6 | 1.6-2.6 | 1.0-2.6 |  |
| Pixel size (Å) | 0.824 | 0.824 | 0.93 | 0.93 | 1.1 | 0.824 |  |
| Initial particle images<br>(no.) | 182,321 | 183,040 | 259,362 | 94,149 | 116,182 | 1,073,937 |  |
| Symmetry imposed | C1 | C1 | C1 | C1 | C1 | Type I<br>C1 | Type II<br>C1 |
| Final particle images<br>(no.) | 182,321 | 68,559 | 38,300 | 10,457 | 15,649 | 102,697 | 3,795 |
| Map resolution (Å) | 3.39 | 2.81 | 3.86 | 3.55 | 3.45 | 2.26 | 3.36 |
| FSC threshold = 0.143 |  |  |  |  |  |  |  |
| Helical rise (Å) | 4.81 | 4.80 | 4.87 | 4.87 | 4.79 | 4.76 | 4.79 |
| Helical twist (°) | -0.56 | -0.63 | -0.60 | -0.62 | -0.64 | -1.18 | -0.77 |
|  | P301L<br>Case 2 |  |  |  |  | P301T<br>Type I | P301T<br>Type II |
| Model Refinement |  |  |  |  |  |  |  |
| Initial model used (PDB) | - |  |  |  |  | - | - |
| Model resolution (Å) | 2.78 |  |  |  |  | 2.23 | 3.00 |
| FSC threshold = 0.5 |  |  |  |  |  |  |  |
| Map sharpening *B* factor<br>(Å²) | -42.2 |  |  |  |  | -36.0 | -12.5 |
| Model composition |  |  |  |  |  |  |  |
| Non-hydrogen atoms | 2,385 |  |  |  |  | 3,232 | 2142 |
| Protein residues | 321 |  |  |  |  | 428 | 288 |
| Ligands | 0 |  |  |  |  | 0 | 0 |
| *B* factors (Å²) |  |  |  |  |  |  |  |
| Protein | 56.4 |  |  |  |  | 41.2 | 79.09 |
| R.m.s. deviations |  |  |  |  |  |  |  |
| Bond lengths (Å) | 0.0061 |  |  |  |  | 0.0040 | 0.003 |
| Bond angles (°) | 1.217 |  |  |  |  | 1.155 | 0.710 |
| Validation |  |  |  |  |  |  |  |
| MolProbity score | 1.38 |  |  |  |  | 1.28 | 1.67 |
| Clashscore | 1.68 |  |  |  |  | 1.51 | 4.80 |
| Poor rotamers (%) | 0.00 |  |  |  |  | 0.00 | 0.00 |
| Ramachandran plot |  |  |  |  |  |  |  |
| Favored (%) | 92.38 |  |  |  |  | 94.29 | 93.62 |
| Allowed (%) | 7.62 |  |  |  |  | 5.72 | 6.38 |
| Disallowed (%) | 0.00 |  |  |  |  | 0.00 | 0.00 |
| PDB | 9GG0 |  |  |  |  | 9GG1 | 9GG6 |
| EMDB | 51319 |  |  |  |  | 51320 | 51325 |

## Notes

### Competing Interest Statement

The authors have declared no competing interest.

